# A base-resolution panorama of the *in vivo* impact of cytosine methylation on transcription factor binding

**DOI:** 10.1101/2021.08.27.457995

**Authors:** Aldo Hernandez-Corchado, Hamed S. Najafabadi

## Abstract

While methylation of CpG dinucleotides is traditionally considered antagonistic to the DNA-binding activity of most transcription factors (TFs), recent *in vitro* studies have revealed a more complex picture, suggesting that over a third of TFs may preferentially bind to methylated sequences. Expanding these *in vitro* observations to *in vivo* TF binding preferences, however, is challenging, as the effect of methylation of individual CpG sites cannot be easily isolated from the confounding effects of DNA accessibility and regional DNA methylation. As a result, the *in vivo* methylation preferences of most TFs remain uncharacterized.

Here, we introduce joint accessibility-methylation-sequence (JAMS) models, which connect the strength of the binding signal observed in ChIP-seq to the DNA accessibility of the binding site, regional methylation level, DNA sequence, and base-resolution cytosine methylation. We show that JAMS models quantitatively explain the TF binding strength, recapitulate cell type-specific TF binding, and have high precision for inferring intra-motif methylation effects. Analysis of 2209 ChIP-seq experiments resulted in high-confidence JAMS models for 260 TFs, revealing that 45% of TFs are inhibited by intra-motif methylation *in vivo*. In contrast, only 16 TFs (6%) preferentially bind to methylated sites, including 11 novel methyl-binding TFs that are mostly from the multi-zinc finger family of TFs.

Our study substantially expands the repertoire of *in vivo* methyl-binding TFs, but also suggests that most TFs that prefer methylated CpGs *in vitro* present themselves as methylation agnostic *in vivo*, potentially due to the balancing effect of competition with other methyl-binding proteins.

## BACKGROUND

Transcription factors (TFs) are key regulators of gene expression. Each TF usually recognizes a specific sequence motif; however, TF binding is affected by several other variables, among which cytosine methylation is traditionally viewed as having a repressive effect on TF binding [1]. However, this traditional view is gradually changing, as more examples are reported of TFs that bind to methylated sequences. These include studies that have reported increased binding of specific TFs to methylated DNA *in vitro* [2], in addition to reports indicating that, for some TFs, a large fraction of their *in vivo* binding sites is highly methylated [3, 4].

While it is tempting to view these anecdotal cases as exceptions rather than a general trend, a recent systematic analysis of TF CpG methylation preferences revealed that, in fact, a large fraction of TFs may bind to methylated CpGs *in vitro*. Based on this study, the effect of methylation is dependent on its position in the binding site, and is heterogeneous within and across TF families [5]. While this study provides *in vitro* evidence for widespread recognition of methylated CpGs by TFs, a comparable systematic analysis of *in vivo* methylation preferences of TFs is still lacking. This is primarily because observing the specific *in vivo* effect of intra-motif CpG methylation is confounded by binding site-specific factors such as DNA accessibility, regional methylation level, and binding site sequence [6–8]. Experimental approaches to control these confounding factors are complicated and resource-exhaustive [9–11], highlighting the need for computational methods to untangle, from these confounding variables, the base-resolution relationship between TF binding affinity and intra-motif CpG methylation.

A few recent studies have proposed computational methods to identify TFs that are affected by CpG methylation *in vitro*. These include efforts to better distinguish bound from unbound sequences using TF binding models that incorporate CpG methylation status [12, 13], as well as tools that expand the ATGC alphabet by adding symbols for methylated cytosines in order to perform methylation-aware *de novo* motif discovery [14, 15]. These methods, however, only report whether methylation improves TF binding prediction without delineating the direction of the effect [13], lack the resolution to investigate the effect of methylation of individual intra-motif cytosines [13], and/or do not consider the confounding effects of DNA accessibility and regional methylation level [12–15]. As a result, even some of the most classic methyl-binding TFs, such as CEBPB [2] and KAISO [16], are not detected by these methods [12].

To overcome these challenges, we introduce Joint Accessibility-Methylation-Sequence models (JAMS), a statistical framework for deconvolving the individual contribution of various factors, including intra-motif CpG methylation, on the *in vivo* strength of TF binding as observed by ChIP-seq. We show that JAMS models are reproducible and generalizable, can capture known CpG methyl-preferences of TFs, and can even predict differential TF binding across cell lines based on changes in intra-motif CpG methylation. Finally, we apply JAMS to a large compendium of ChIP-seq experiments to systematically explore the CpG methylation preferences of TFs across different families.

## RESULTS

### Modeling the joint effect of accessibility, methylation and sequence on TF binding

Several factors work together to determine the TF binding strength for a specific binding site. First, the sequence of the binding site determines the TF affinity, given that the majority of TFs are sequence-specific. Secondly, for most TFs, the existing level of DNA accessibility heavily influences TF binding [7, 8]. Thirdly, regional methylation outside the TFBS may affect the TF binding strength, for example by recruiting Methyl-CpG-binding domain (MBD) proteins, which in turn recruit chromatin remodelers [6]. Therefore, in order to examine the specific effect of methylation of the TFBS on TF binding affinity, we need to jointly model it together with these confounding factors.

For this purpose, we developed Joint Accessibility-Methylation-Sequence models (JAMS), which quantitatively explain both the pull-down and background signal in ChIP-seq experiments (https://github.com/csglab/JAMS). The JAMS model for each ChIP-seq experiment considers the pull-down read density as a combination of a background signal and a TF-specific signal. On the other hand, the read count profiles obtained from control experiments (e.g. input DNA) purely reflect the background signal (**Fig. 1A**). Each of the background and TF-specific signals, in turn, is modeled as a function of the peak sequence, chromatin accessibility profile along the peak, regional methylation level, and base-resolution intra-motif CpG methylation (**Fig. 1B-C**). JAMS converts these associations into a generalized linear model, whose parameters can be inferred by fitting simultaneously to both pull-down and control read counts. To ensure that JAMS can correctly learn the features associated with both TF-specific and background signals, we fit the model to the read counts across peaks with a wide range of pulldown-to-control signal ratio. These include not only the peaks that have significantly high pull-down signal, but also peaks with low pull-down signal as well as genomic locations with significantly high background signal. For model fitting, an appropriate error model is needed that connects the expected (predicted) signal at each peak to the observed read counts—we use negative binomial with a log-link function in this work (**Fig. 1D**; see **Methods** for details).

**Figure 1.**
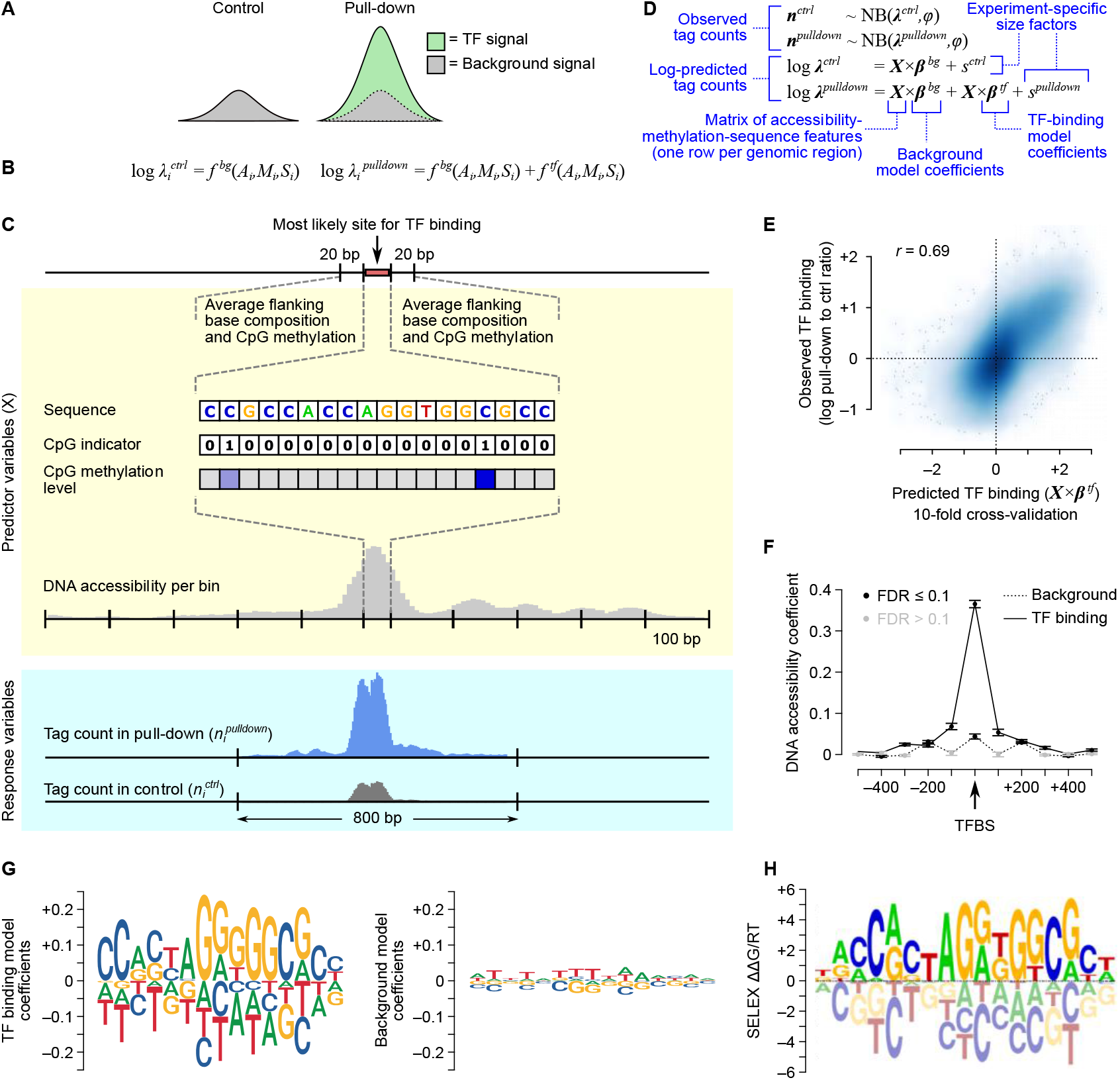
Overview of JAMS model. (**A**) At each genomic region *i*, the JAMS model considers the control tag count (left) or the pull-down tag count (right) as a combination of background and/or TF-binding signals at that position. (**B**) Each of these signals are then modeled as a function of accessibility (*A_i_*), methylation (*M_i_*), and sequence (*S_i_*) at each region *i*. (**C**) Schematic summary of the predictor features extracted for each genomic location and the outcome variables. (**D**) The specifications of the generalized linear model used by JAMS. (**E**) Comparison between the observed and predicted CTCF binding signal in HEK293 cells [20]. (**F**) DNA accessibility coefficients learned by the CTCF JAMS model; each dot corresponds to the effect of accessibility at a 100bp-bin. (**G**) Sequence motif logos representing the TF binding specificity learned by JAMS (left) and the effect of sequence on the background signal (right). JAMS motif logos are plotted using ggseqLogo [23], with letter heights representing model coefficients. (**H**) The known CTCF binding preference (based on SELEX [24]); SELEX motif logo was obtained from the CIS-BP database [25].

In order to examine the ability of JAMS models to recover the *in vivo* binding preferences of TFs, we first applied it to ChIP-seq data from CTCF, a widely studied TF that is constitutively expressed across cell lines and tissues [17, 18] and has a long residence time on DNA [19]. We initially focused on the cell line HEK293, and generated a JAMS model of CTCF binding in this cell line using previously published ChIP-seq [20], WGBS [21], and chromatin accessibility data [22] (**Methods**). To evaluate the performance of the JAMS model, we used 10-fold cross-validation, and examined the correlation between the predicted TF-specific signal and the observed pulldown-to-control signal ratio across the peak regions. As **Fig. 1E** shows, the JAMS model predictions correlate strongly with the pulldown-to-control signal ratio (Pearson *r*=0.69), suggesting that accessibility-methylation-sequence features can quantitatively predict the CTCF-binding strength.

Examining the coefficients of the fitted JAMS model, we observed that DNA accessibility, especially at the peak center, has a strong effect on the TF-specific signal (which only affects the pull-down read count), but limited effect on the background ChIP-seq signal (which affects both the control and pull-down read counts; **Fig. 1F**). Nonetheless, the effect on background signal was still statistically significant, consistent with previously observed bias of DNA sonication toward accessible chromatin regions [26]. Importantly, sequence features at the TF binding site are strongly predictive of the CTCF binding strength, while they have limited and diffuse effect on the background signal (**Fig. 1G**). The sequence model learned by JAMS is highly correlated with the known motif for CTCF (*r*=0.86, **Fig. 1H**), suggesting that JAMS models can recapitulate the underlying biology of TF binding.

### JAMS models reveal the contribution of CpG methylation to TF binding

By jointly considering the contribution of accessibility, methylation and sequence to TF binding, JAMS models should be able to deconvolve the specific effect of methylation from the confounding effect of other variables. To begin to explore this possibility, we examined the JAMS model of CTCF. For this purpose, in addition to the widely used sequence motif logos, we developed “dot plot logos” to enable easier visual inspection of JAMS coefficients that correspond to sequence and methylation effects. As **Fig. 2A** shows, the JAMS model of CTCF binding in HEK293 cells suggests that methylation of C2pG3 and C12pG13 of the binding site has a significantly negative effect on CTCF binding (but not on the background signal; **Supplementary Fig. 1**). In other words, while a large fraction of CTCF binding sites have CpGs at positions 2/3 and 12/13, CTCF preferentially binds when these CpGs are not methylated.

**Figure 2.**
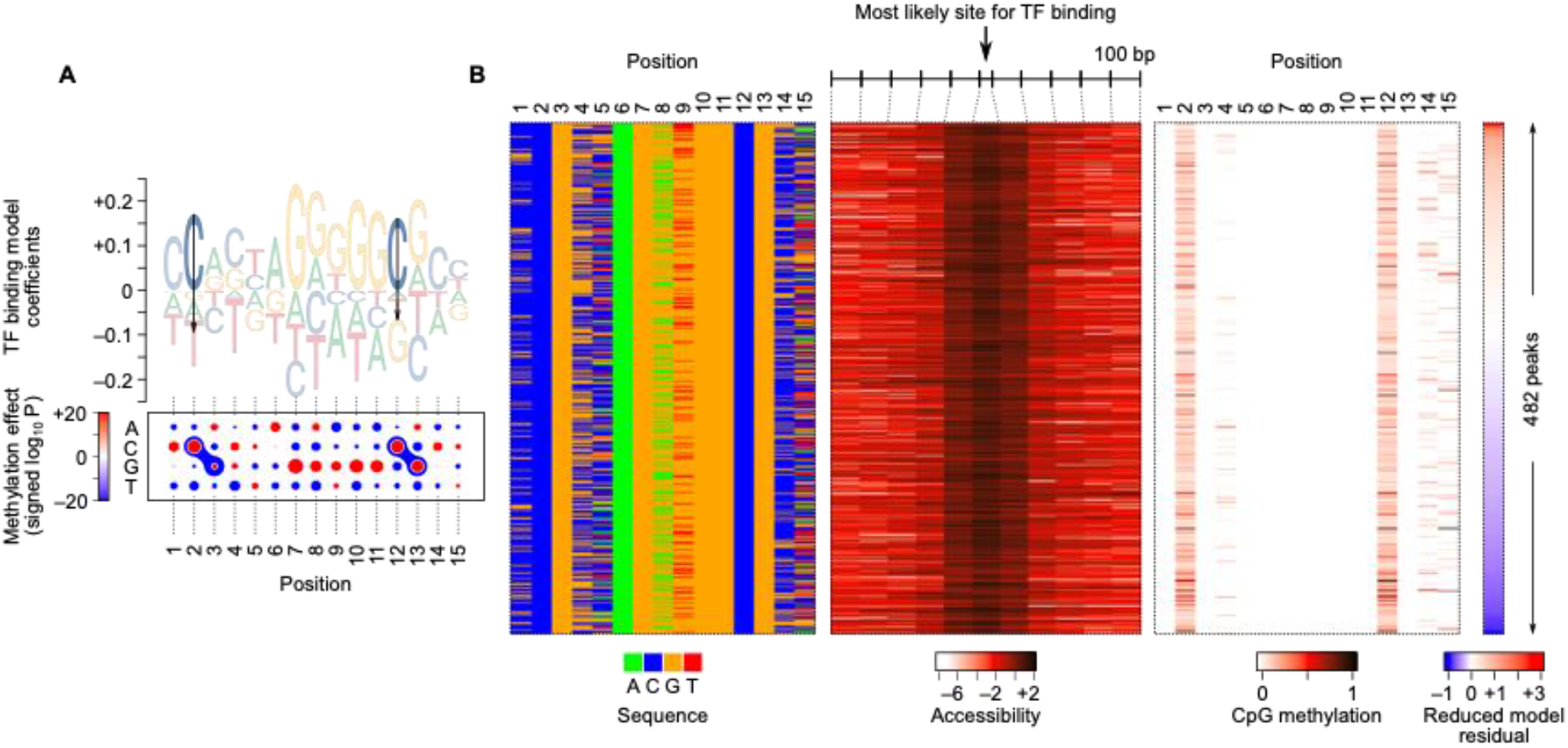
CpG methylation preference of CTCF in HEK293 cells. (**A**) Motif logo and dot plot representations of the sequence/methylation preference of CTCF. The logo (top) shows methylation coefficients as arrows, with the arrow length proportional to the mean estimate of methylation effect. The dot plot (bottom) shows the magnitude of the preference for each nucleotide at each position using the size of the dots, with red and blue representing positive and negative coefficients, respectively. The dumbbell-like shapes demarcate the CpG dinucleotides with significant methylation effects (at FDR<1×10^-5^). The color of the dumbbell shows the signed logarithm of P-value of the methylation coefficient, with red and blue corresponding to increased or decreased binding to methylated C, respectively. (**B**) Heatmap representation of the sequence (left), accessibility (middle), and CpG methylation (right), for a subset of CTCF peaks that have high DNA accessibility, a close sequence match to the initial CTCF motif, and CpGs at dinucleotide positions 2/3 and 12/13. Peaks (rows) are sorted by the residual of a reduced JAMS model that does not use the methylation level of C2pG3 and C12pG13 for predicting the CTCF binding signal. Note that in the methylation heatmap (right), the methylation level of a CpG dinucleotide is shown in the column that corresponds to the position of the C nucleotide. For example, values in column 2 correspond to the methylation level of the C2pG3 dinucleotide.

To ensure that this observation is not confounded by other variables such as accessibility and the average local methylation level, we also trained JAMS models with all the variables except the CpG methylation level at each binding site position; we then compared these reduced models to the full model using a likelihood ratio test. This analysis revealed that removing the information about methylation levels of C2pG3 or C12pG13 significantly reduces the fit of the model to the observed data (**Supplementary Fig. 2**). Therefore, the CpG methylation level in these positions is informative about CTCF binding signal even after considering the effect of other confounding variables such as sequence, accessibility, and the average methylation of flanking regions. The independent effect of CpG methylation on CTCF binding can also be observed after stratification of CTCF peaks based on the confounding variables. Specifically, we repeated the JAMS modeling after removing the variables that represent the TF-specific contribution of methylation at dinucleotides C2pG3 and C12pG13, and sorted the peaks by the residual of this model (i.e. by the ChIP-seq signal that could not be explained by the reduced model). As **Fig. 2B** shows, even if we focus on the peaks with similar DNA sequence and accessibility, the residual of the reduced model still correlates negatively with CpG methylation at positions 2/3 and 12/13. In other words, peaks whose signal is smaller than what the reduced model predicts have higher CpG methylation, supporting the negative effect of CpG methylation on CTCF binding. Importantly, our observation that CpG methylation negatively affects CTCF binding is consistent with previous reports on CTCF methylation preferences *in vivo* [27] and *in vitro* [28]. Our results are also reproducible across different cell lines, as we obtained similar JAMS models using CTCF ChIP-seq, WGBS, and accessibility data from several other cell lines (**Supplementary Fig. 3**). These results overall suggest that JAMS models have the potential to faithfully recapitulate the methylation preferences of TFs using ChIP-seq data.

### Differential TF binding across cell lines can be explained using JAMS models

A model that encodes the intrinsic binding preference of a TF should be able to predict the ChIP-seq signal of that TF in new contexts, such as in previously unseen cell types that were not used in model training. We began to examine this possibility by investigating the transferability of the CTCF model that was learned in HEK293 cells to other cell types. We used DNase-seq and WGBS data (**Methods**) from six cell lines (H1, GM12878, HeLa-S3, HepG2, and K562) to predict the CTCF binding signal (using the HEK293-trained JAMS model), and compared the predictions to experimental CTCF ChIP-seq data obtained for each cell type. We observed that the CTCF JAMS model that was trained on HEK293 data could successfully predict the ChIP-seq pulldown-to-control ratio in other cell types, with a performance comparable to JAMS models that were specifically trained on the data from each type (**Table 1**). These results support the transferability of JAMS models across cell types.

**Table 1:** Pearson correlation (*r*) between observed and predicted CTCF-binding across cell types. The third column shows *r* between observed and cross-validated JAMS predictions for models that were trained on each individual cell type. The fourth column shows the *r* between the predictions of the JAMS model that was trained on HEK293 and the observed ChIP-seq data in other cell lines.

| Cell line | ChIP-seq peaks ($n$) | 10-fold CV | HEK293-trained $r$ |
| --- | --- | --- | --- |
| HEK293 | 135,717 | 0.69 | - |
| H1 | 128,123 | 0.72 | 0.62 |
| GM12878 | 39,535 | 0.69 | 0.54 |
| HeLa-S3 | 65,865 | 0.72 | 0.60 |
| HepG2 | 81,188 | 0.73 | 0.64 |
| K562 | 85,122 | 0.74 | 0.68 |

The above analysis shows that the JAMS models learned from one cell type can be transferred to another cell type. However, the majority of CTCF binding sites are shared across different cell types; therefore, it is not immediately clear to what extent this transferability corresponds to cell-invariant features of the JAMS model (sequence) as opposed to potentially cell type-specific features (methylation and accessibility). In fact, one of the most challenging aspects of modeling TF binding is the ability to identify TF binding sites that are differentially occupied across cell types [29]. To understand the extent to which differential accessibility and methylation of DNA drives differential CTCF binding, and the extent to which these effects can be captured by JAMS, we decided to use the JAMS model learned from HEK293 cells to predict differential binding of CTCF in other cell lines. We started by identification of differentially bound CTCF peaks in pairwise comparisons of cell lines listed in **Table 1**. For any given two cell lines, we used the log-fold change (log-fc) in the pulldown-to-control ratio as the measure of differential binding (**Fig. 3A**). The mean and standard error of mean (SEM) of this metric was calculated using a statistical model that assumes a negative binomial distribution for the tag counts, which also allows us to calculate a P-value for the null hypothesis that log-fc is equal to zero (see **Methods**). Application of this method to all pairwise cell comparisons revealed the largest number of statistically significant differential CTCF peaks between GM12878 and HeLa-S3 cells (**Fig. 3B**); therefore, we focused on prediction of the differential peaks between these two cell lines using the HEK293 JAMS model of CTCF. Specifically, we used the JAMS model to predict the CTCF binding signal in each of the GM12878 and HeLa-S3 cell lines (based on the accessibility and methylation data of each cell line), and then calculated the difference of the JAMS predictions (in log-scale) between the two cells. As shown in **Fig. 3C**, the JAMS-predicted changes in CTCF binding are strongly correlated with the experimental logFC values (*r*=0.40 across peaks with logFC standard error of mean <1.28; see **Supplementary Fig. 4** for details on the choice of cutoff). These results suggest that the CTCF JAMS model can quantitatively predict the change in CTCF binding strength based on differential accessibility and methylation. Importantly, for the set of peaks that pass the statistical significance threshold for differential binding between the two cell lines (FDR<0.1), the correlation between JAMS predictions and experimental log-fc reaches as high as 0.84 (**Fig. 3C**), with JAMS being able to distinguish GM12878-specific from HeLa-S3-specific binding events with 95% accuracy.

**Figure 3.**
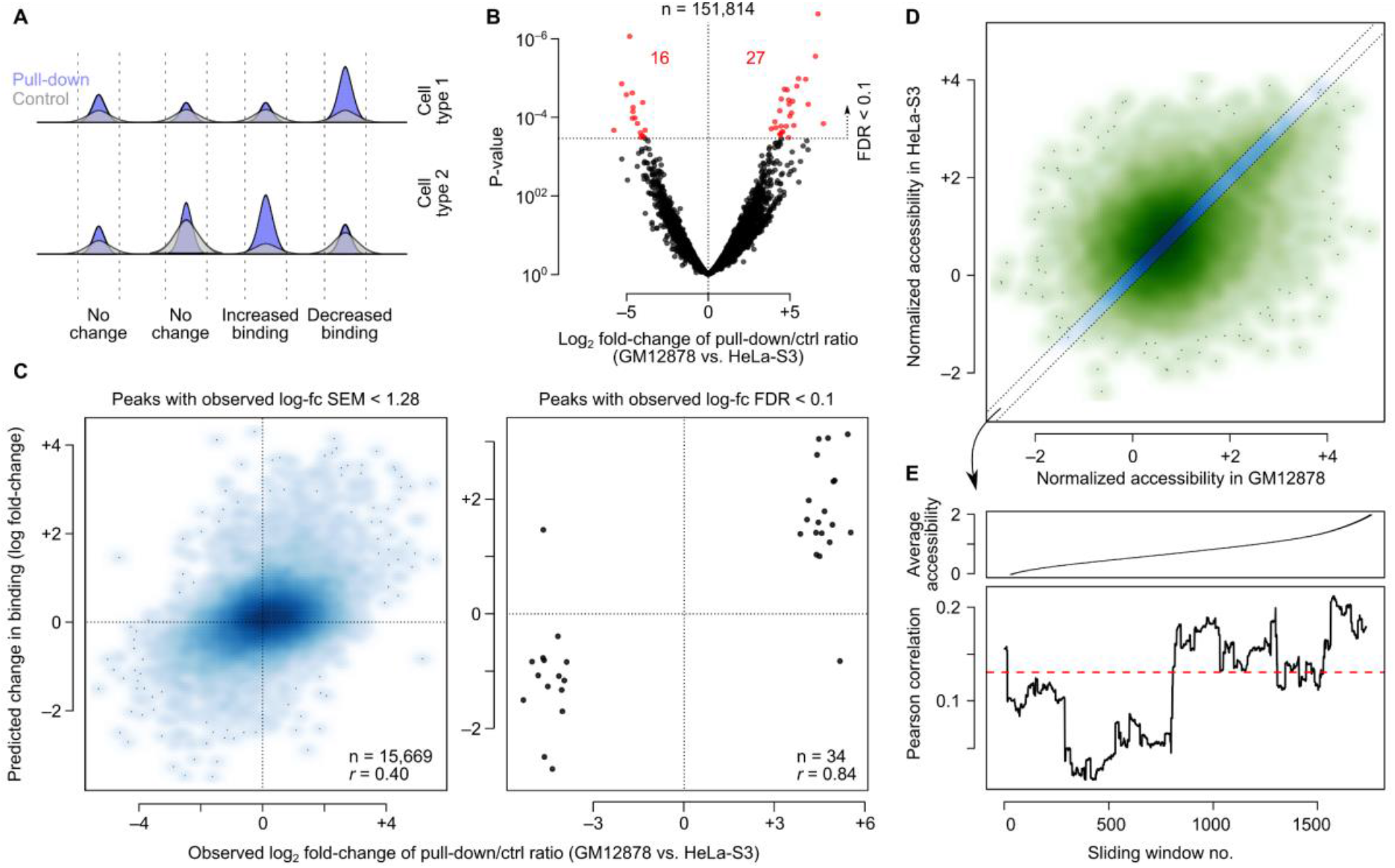
Prediction of differentially bound CTCF peaks using JAMS. (**A**) Schematic representation of identifying differentially bound peaks based on the combination of pulldown and control signal in two cell lines. See **Methods** for details. (**B**) Volcano plot showing differential binding of ChIP-seq peaks between GM12878 and HeLa-S3. Significant peaks at FDR < 0.1 are shown in red. (**C**) Left: Scatter plot of JAMS-predicted changes in CTCF binding and observed differential binding between GM12878 and HeLa-S3 cells. Peaks with observed logFC SEM <1.3 are included. Right: Limited to peaks that pass FDR<0.1 for differential binding of CTCF. (**D**) Comparison of the accessibility of putative CTCF peaks between two cell lines. The diagonal band in the middle (blue) shows the region that was selected as no-change in accessibility (difference in accessibility < 0.2). (**E**) Predicting differential CTCF binding for peaks with no change in accessibility. Peaks were ranked by accessibility, and the correlation between predicted and observed logFC of CTCF binding was calculated for sliding windows of 500 peaks (bottom). The average accessibility for each sliding window is shown on top.

We note that many of the CTCF binding sites are differentially accessible between GM12878 and HeLa-S3 (**Fig. 3D**), which may drive the differential binding. To specifically examine the role of differential methylation in driving cell type-specific CTCF binding, we further limited our analysis to the set of peaks that had similar accessibility in both cell lines (**Fig. 3D**), and also removed all the JAMS predictor variables corresponding to accessibility. We observed that this reduced JAMS model can still predict differential CTCF binding among the peaks that are not differentially accessible (*r*=0.14 between predicted and observed log-fc across n=2232 peaks, p-value < 2×10^-11^; **Fig. 3E**). This correlation increases to 0.22 for the set of peaks that have high accessibility in both cell lines (**Fig. 3E**), suggesting that the effect of differential CpG methylation is most noticeable when the putative CTCF binding site is accessible in both cell lines.

Overall, these analyses suggest that JAMS models can predict differential TF binding across cell types, including differential TF binding events that are driven by changes in the methylation of the putative binding sites. The ability of JAMS to predict cell type-specific TF binding events further highlight its reliability in capturing the determinants of TF binding using ChlP-seq data.

### Systematic inference of the *in vivo* methyl-binding preferences of 260 TFs using JAMS

To identify TFs whose *in vivo* binding is positively or negatively affected by methylation of intra-motif CpGs, we decided to apply JAMS to a comprehensive compendium of ChIP-seq data for a wide range of TFs. We collected and uniformly processed data from 2209 ChIP-seq and ChIP-exo experiments [20, 22, 30], covering the *in vivo* binding profiles of 604 TFs in six cell lines (**Supplementary Table 1**), along with the WGBS and DNase-seq assays in those cell lines (**Supplementary Table 2**). On average, we identified ~60k peaks per ChIP-seq experiment using the permissive P-value threshold of 0.01 (**Supplementary Fig. 5**). We then used the peak tag counts to fit a JAMS model to each ChIP-seq experiment. We noticed that the quality of the JAMS models, measured by the Pearson correlation between the predicted and observed TF-specific signal, varied substantially across the experiments, with correlations ranging from 0 to 0.8 (median 0.48; **Supplementary Fig. 5**). This variation may reflect a multitude of factors, including the ChIP-seq data quality as well as the extent to which the TF signal can be explained by our model specifications. We therefore decided to keep only a subset of high-confidence models. Specifically, we selected at most one representative model per TF based on the following criteria: (i) the model should have used at least 10,000 peaks for training, (ii) Pearson correlation >0.2 between the predicted and observed TF-specific signal after cross-validation, (iii) Pearson correlation >0.3 between the known and JAMS-inferred sequence motif, (iv) and low contribution of the sequence to the background signal compared to the TF-specific signal (control-to-pulldown ratio of the sequence coefficients mean < 0.4). As an example, in **Supplementary Fig. 6** we show two JAMS models for BHLHE40, obtained from two different ChIP-seq experiments, only one of which passes all the criteria mentioned above. Overall, we obtained high-confidence JAMS models for 260 TFs, spanning a range of TF families (**Fig. 4D** and **Supplementary Table 3**).

**Figure 4.**
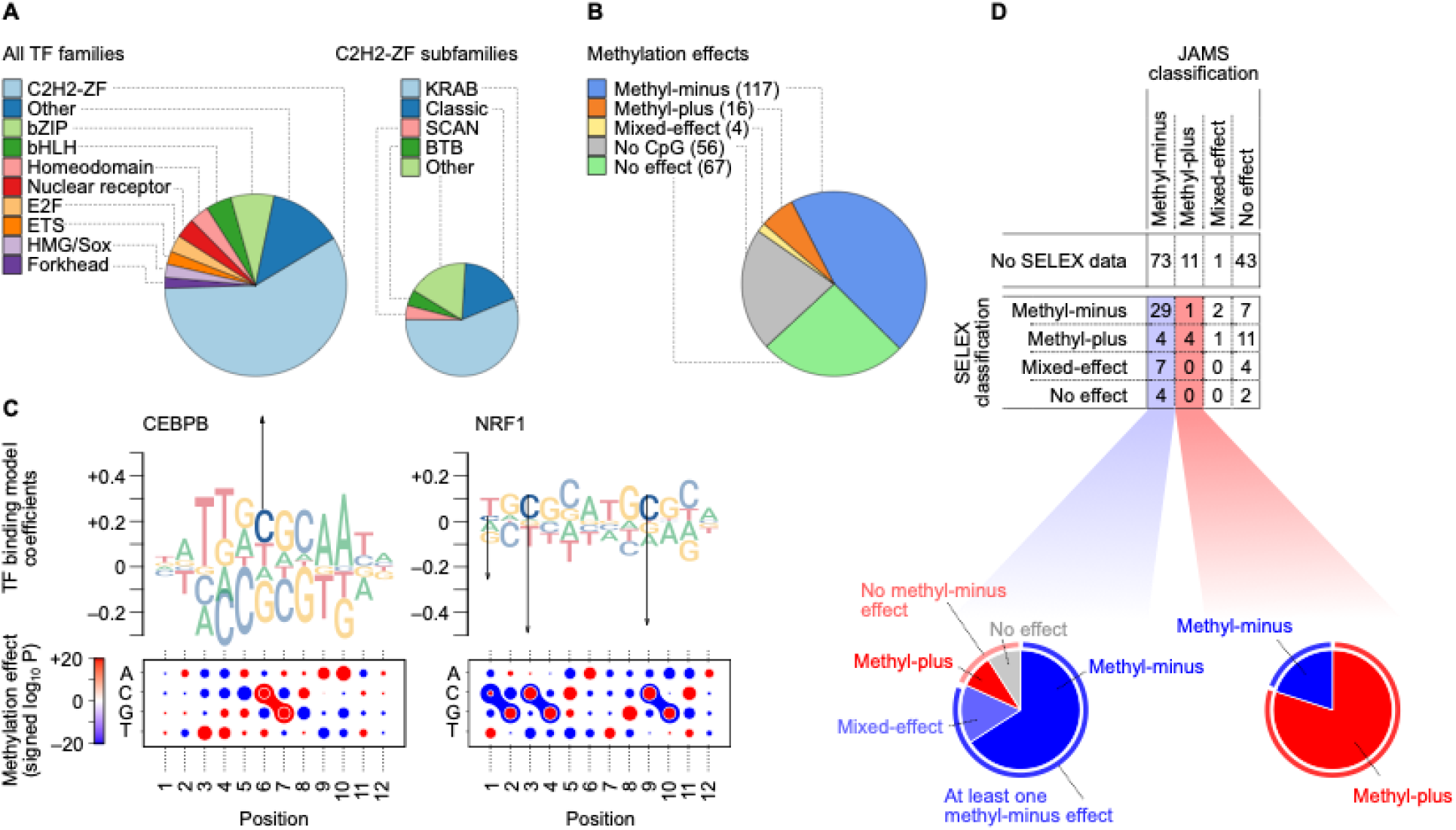
Systematic application of JAMS. (**A**) Pie charts of the main TF families (left) and C2H2-ZF proteins subfamilies (right) for TFs with at least one high-quality JAMS model. (**B**) Pie chart of the methyl-binding preferences of TFs with at least one high quality JAMS model. We obtained high-quality models for a total of 260 TFs. (**C**) Motif logos and dot plot representations of the sequence/methylation preference of CEBPB (left) and NRF1 (right), as inferred by JAMS (see **Fig. 2A** for a description of how these representations should be interpreted). (**D**) The table on the top shows the confusion matrix of TF classifications by JAMS (columns) and methyl/bisulfite-SELEX [5] (rows). The pie charts at the bottom illustrate how JAMS methyl-minus (left) and methyl-plus (right) predictions correspond to different SELEX-based classifications.

After selecting one JAMS model per TF, we used the JAMS-inferred effects of methylation to classify the TFs according to their inferred methyl-binding preferences. We use a notation similar to Yin et al. [5]. Specifically, we classified a TF as (a) methyl-minus if its JAMS model included at least one significantly negative mCpG effect (FDR<1×10^-5^), (b) methyl-plus if the model included at least one significantly positive mCpG effect, (c) mixed-effect if the model included both significantly positive and negative mCpG effects, (d) and no-effect if the motif included a CpG but its methylation level did not have a significant effect. Overall, we found 117 methyl-minus TFs, 16 methyl-plus TFs, four mixed-effect TFs, and 67 TFs with no significant mCpG effects; we also identified a set of 56 TFs without a CpG site in their binding site (**Fig. 4B**).

To understand whether our JAMS-based classification captures known methyl-binding preferences of TFs, we started by examining a few TFs whose methyl-binding preferences have been extensively studied *in vitro* and *in vivo*, including CEBPB and NRF1. Using protein-binding microarrays (PBMs), Mann et al. have previously reported enhanced binding of CEBPB to its CpG-containing target sequence when the array probes were methylated [2], consistent with the observation that a large fraction of the genomic binding sites of CEBPB is highly methylated *in vivo* [3]. The JAMS model for CEBPB (**Fig. 4C** and **Supplementary Fig. 7A-D**) is concordant with these previous reports, showing that methylation of C6pG7 dinucleotide in the CEBPB target sequence has a positive effect on its binding strength. This effect is in fact highly reproducible, and is present in three out of four JAMS models that we obtained using different CEBPB ChIP-seq experiments. Another well studied TF is NRF1, which has been found to be sensitive to CpG methylation of DNaseI-hypersensitive sites in murine stem cells [10]. Moreover, Cusack et al. found that NRF1 preferentially binds to unmethylated DNA even after accounting for changes in DNA accessibility caused by the recruitment of HDACs to methylated CpGs through MBD proteins [9]. Consistent with these reports, we found that methylation of C3pG4 and C9pG10 dinucleotides in the NFR1 target sequence has a negative effect on its binding (**Fig 4C and Supplementary Fig. 7E-H**); these effects were consistent across all the cell lines we analyzed.

The above examples suggest that JAMS models are consistent with previously reported methylation preferences of TFs. However, there are only a handful of TFs whose methylation preferences have been validated *in vivo*. Therefore, to systematically evaluate our JAMS-based classification of TFs, we compared our inferred methyl-binding preferences with *in vitro* preferences obtained using methyl-SELEX and/or bisulfite-SELEX [5]. Overall, 76 out of the 260 TFs that we studied here have methyl/bisulfite-SELEX data (**Fig. 4D**). These included 44 TFs that we classified as methyl-minus based on *in vivo* data; 29 of these TFs (~66%) were also identified as methyl-minus by SELEX, and another 7 TFs (16%) were identified as mixed-effect. This suggests that our approach has ~82% precision for identification of TFs that are negatively affected by CpG methylation in at least one position in their target sequence. On the other hand, out of 39 methyl-minus TFs found by SELEX, 31 were also classified as either methyl-minus or mixed-effect by JAMS, suggesting that ~79% of *in vitro*-observed methyl-minus effects can be captured using *in vivo* data.

Similarly, out of five JAMS-based methyl-plus TFs that have bisulfite-SELEX data [5], four were classified as methyl-plus based on SELEX (**Fig. 4D**), suggesting a precision of ~80%. However, despite this high precision, only 5 out of 20 SELEX-based methyl-plus TFs are identified as either methyl-plus or mixed-effect by JAMS—this suggests that a relatively small fraction of *in vitro* methyl-plus effects can also be observed *in vivo*. Nonetheless, we found 11 methyl-plus TFs that were previously unclassified—this is in addition to 73 previously unclassified methyl-minus and one novel mixed-effect TF, highlighting the ability of JAMS models in revealing novel TF methyl preferences.

**Fig. 5A** shows the distribution of different methyl-preferences across main TF families. We noticed that a disproportionately large number of methyl-plus TFs belong to the C2H2-ZF family (methyl-preferences of these TFs are shown in **Fig. 5B**). More specifically, among the KRAB domain-containing members of the C2H2-ZF family whose binding is significantly affected by methylation, ~24% preferentially bind to methylated CpGs (**Table 2**), compared to only ~12% of non-KRAB TFs (Fisher’s exact test P<0.009, **Supplementary Table 4**). This is an intriguing observation, given that a majority of KRAB-ZF proteins evolved to specifically bind and repress transposable elements, which largely reside in highly methylated genomic regions[31]. It is notable that we observed this methyl-plus effect even though we removed all repetitive genomic regions from our analysis (see **Methods**). Our observation suggests that many of the KRAB-ZF proteins preferentially bind to methylated instances of their target sequence, potentially allowing them to distinguish the transposable elements from other genomic regions that contain their preferred binding sequence. In fact, ~56% of all methyl-plus TFs that we identified are KRAB-ZF proteins, suggesting that recognition of methylated transposable elements might have been a primary force in the evolution of methyl-binding TFs.

**Figure 5.**
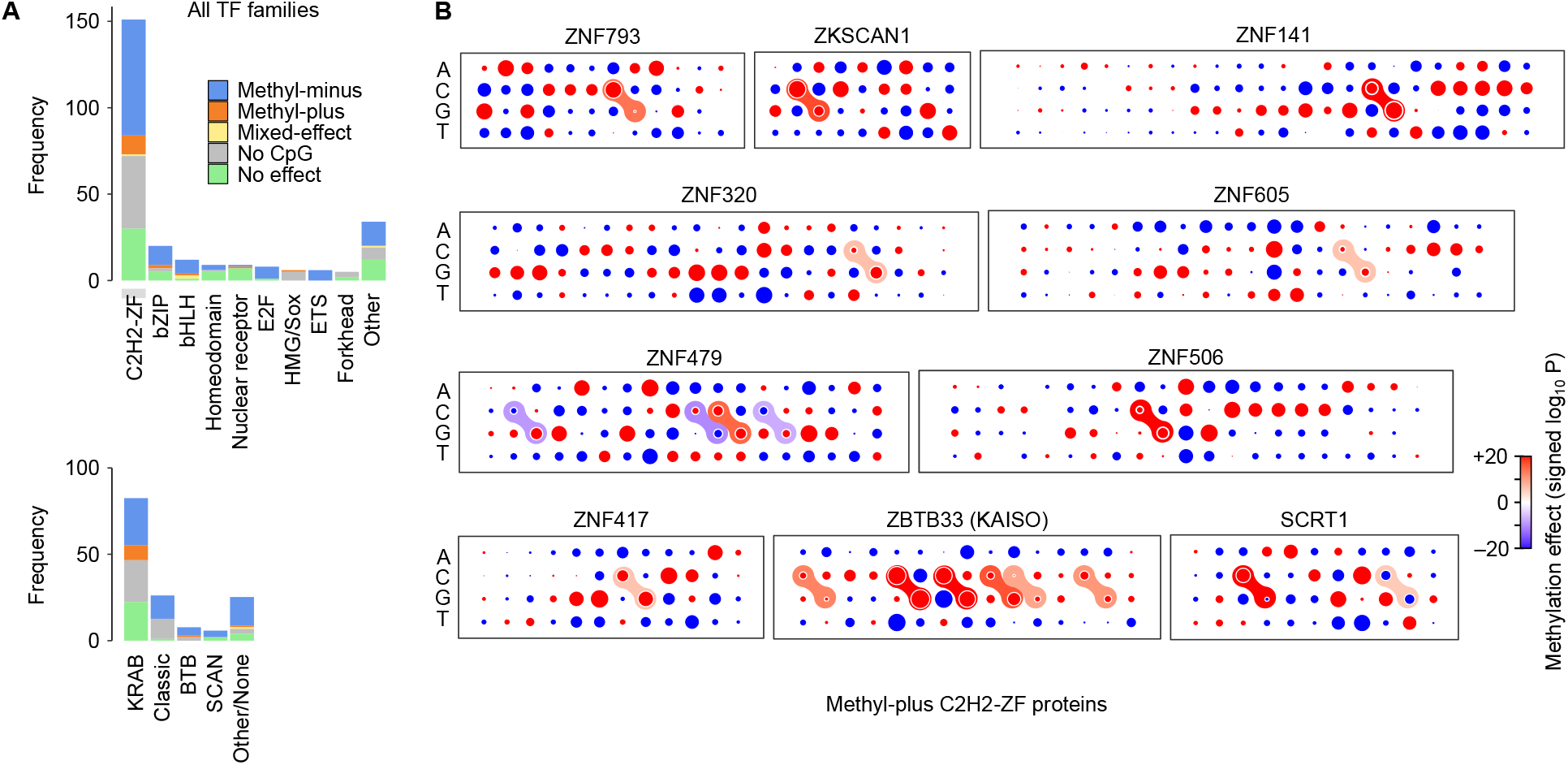
Methylation preferences per TF family. (**A**) Top: Stacked bar plots showing the distribution of TF methylation preferences inferred with JAMS, grouped by TF families. Bottom: The distribution of methylation preferences for C2H2-ZFP subfamilies. (**B**) Dot plot representation of the methylation preferences for the C2H2-ZF proteins that, based on JAMS analysis, are methyl-plus. See **Supplementary Fig. 8** for motif logos.

**Table 2.** TFs with methyl-plus and mixed-effect methyl-binding preferences, as inferred by JAMS using *in vivo* data. For mixed-effect TFs, both the position at which a positive methylation effect was observed as well as the position with a negative methylation effect are indicated. See **Supplementary Fig. 8** for motif logos.

| Protein | Family | JAMS call | Effect of methylation by position |  | SELEX call [5] |
| --- | --- | --- | --- | --- | --- |
|  |  |  | Positive | Negative |  |
| CEBPB | bZIP | Methyl-plus | 6 |  | Methyl-plus |
| SCRT1 | C2H2 ZF | Methyl-plus | 3 |  | Methyl-plus |
| CEBPG | bZIP | Methyl-plus | 6 |  | Methyl-plus |
| ZBTB33 (KAISO) | C2H2 ZF (BTB) | Methyl-plus | 5, 7 |  | Methyl-plus |
| TCF7 | HMG/Sox | Methyl-plus | 2 |  | Methyl-minus |
| ZKSCAN1 | C2H2 ZF (KRAB+SCAN) | Methyl-plus | 2 |  |  |
| ZNF793 | C2H2 ZF (KRAB) | Methyl-plus | 7 |  |  |
| ZNF141 | C2H2 ZF (KRAB) | Methyl-plus | 17 |  |  |
| ZNF320 | C2H2 ZF (KRAB) | Methyl-plus | 17 |  |  |
| ZNF605 | C2H2 ZF (KRAB) | Methyl-plus | 15 |  |  |
| ZNF479 | C2H2 ZF (KRAB) | Methyl-plus | 11 |  |  |
| ZNF490 | C2H2 ZF (KRAB) | Methyl-plus | 7 |  |  |
| ZNF506 | C2H2 ZF (KRAB) | Methyl-plus | 5 |  |  |
| ZNF417 | C2H2 ZF (KRAB) | Methyl-plus | 16 |  |  |
| NR2F2 | Nuclear receptor | Methyl-plus | 5, 8 |  |  |
| TFAP4 | bHLH | Methyl-plus | 7 |  |  |
| SP1 | C2H2 ZF | Mixed-effect | 5 | 8 | Methyl-plus |
| USF1 | bHLH | Mixed-effect | 7 | 5 | Methyl-minus |
| USF2 | bHLH | Mixed-effect | 7 | 5 | Methyl-minus |
| NFYB | NFYB/HAP3 | Mixed-effect | 9 | 13 |  |

## DISCUSSION

In this study, we built Joint Accessibility-Methylation-Sequence (JAMS) models to capture the relationship between TF binding and DNA methylation *in vivo*. Our approach models the TF binding as a function of DNA accessibility, sequence and methylation at and around TF binding sites, while separating the background from TF-specific signals. By systematic application of JAMS to a large compendium of ChIP-seq datasets and comparison to SELEX-based *in vitro* data [5], we showed the reliability of methylation preferences identified by JAMS, with ~80% of methyl-plus and methyl-minus TFs found by JAMS showing a concordant effect *in vitro*. In addition, we characterized the methylation preferences of 128 TFs that were not previously studied by bisulfite- or methyl-SELEX, revealing 73 novel methyl-minus and 11 novel methyl-plus TFs (**Fig. 4D**).

An intriguing observation from the comparison of *in vivo* JAMS models and *in vitro* SELEX models (**Fig. 4D**) is that the methyl-binding capacity of TFs overall decreases *in vivo* compared to *in vitro*: Most TFs that are methyl-plus *in vitro* become indifferent to the methylation status of CpGs *in vivo* (11 out of 20) or even become methyl-minus (4 out of 20); most TFs that are indifferent to methylation *in vitro* become methyl-minus *in vivo* (4 out of 6); and most TFs that are methyl-minus *in vitro* also present themselves as methyl-minus *in vivo* (29 out of 39). This trend can even be seen at the level of individual binding site positions; for example, while *in vitro* studies have found that CTCF binding is sensitive to methylation of the dinucleotide C2pG3 of its binding sequence [28], we found that methylation of C12pG13 has an additional negative effect on CTCF binding *in vivo*, which is not observed *in vitro*.

One possible explanation for this shift toward methylation avoidance is the direct competition of TFs with MBD proteins. While JAMS is able to capture the indirect effect of MBD proteins on DNA accessibility (through recruitment of chromatin modifiers), as well as potential MBD recruitment through flanking mCpGs, it currently does not model the direct competition of TFs and MBD proteins for binding to intra-motif mCpG sites. This undetected direct competition could affect the interpretation of our model parameters: methylation coefficients obtained by JAMS models should be more accurately interpreted as the affinity of a TF toward mCpG sites “relative” to the affinity of other competing factors, such as MBD proteins. **Fig. 6** schematically shows the most common scenarios that may arise from this competition and their estimated frequency based on our JAMS-SELEX comparison.

**Figure 6.**
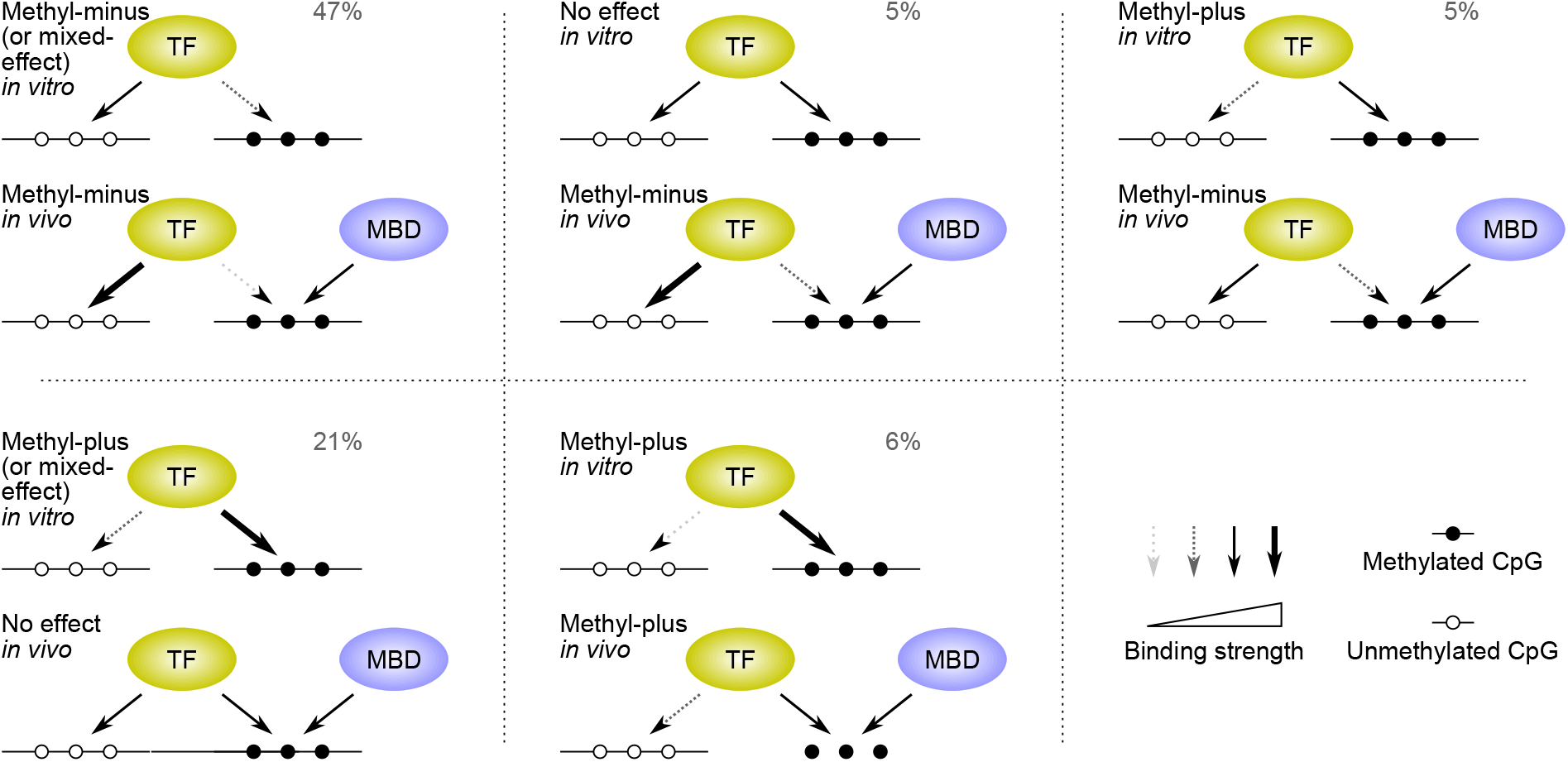
Schematic presentation of how competition with MBD proteins may affect TF binding. Each panel shows how *in vitro*-observed mCpG preferences may present themselves *in vivo* in the presence of competing mCpG-binding proteins such as MBDs. The percentages indicate the estimated frequency of each scenario among CpG-binding TFs. For example, 80% (4 out of 5) of JAMS-based methyl-plus TFs that have SELEX data show methyl-plus preference *in vitro*, and a total of 16 methyl-plus TFs are identified by JAMS. Therefore, ~13 out of these 16 TFs are expected to be *in vitro* methyl-plus TFs that remain methyl-plus *in vivo*, corresponding to ~6% (13 out of 204) of all CpG-binding TFs.

Accordingly, for the majority of *in vitro* methyl-plus TFs, their competition with MBD proteins leads to their apparent indifference to methylation *in vivo*, resulting in equal recognition of methylated and unmethylated CpGs by these TFs—we have identified a total of 67 *in vivo* methyl-indifferent TFs, ~60% of which is expected to show some degree of mCpG preference in the absence of MBD proteins *in vitro*. On the other hand, only the TFs with the strongest affinity toward methylated CpGs are expected to outcompete MBD proteins and bind preferentially to mCpG sites *in vivo*—our analysis has identified 16 such TFs (**Table 2**), including 11 novel methyl-plus TFs, most of which belong to the C2H2-ZF class of proteins.

These results represent, to our knowledge, the largest resource for exploring the *in vivo* effect of methylation on TF binding. They suggest that preferential binding of TFs to *in vivo* methylated CpGs is not rare, but also not as pervasive as it may appear from *in vitro* experiments. Instead, TF affinity for mCpGs often equilibrates with the mCpG-binding activity of other proteins such as MBDs, resulting in the overall methylation-agnostic activity of ~20% of CpG-binding TFs.

## METHODS

### Methods overview

To understand the relationship between DNA methylation and TF binding, we began by retrieving and analyzing WGBS, ChIP-seq, and DNase-seq data from different TFs in several cell lines. We developed a method to jointly model these data sets to predict TF-specific binding, and benchmarked it on CTCF ChIP-seq data in HEK293 cells. We expanded our CTCF studies by obtaining differential binding sites of CTCF between different cell lines, and examined whether, using our method, we can predict differential binding that was caused by DNA methylation changes. Finally, we applied our method to a comprehensive collection of ChIP-seq data to systematically study the *in vivo* effect of DNA methylation on TF binding.

### ChIP-seq data processing, peak calling, and peak signal quantification

We limited our analysis to ChIP-seq experiments performed in HepG2, K562, HEK293, GM12878 and HeLa-S3 cell lines, given the availability of high-depth WGBS and DNase-seq data for these cell lines. ChIP-seq and ChIP-exo raw reads were retrieved from four main sources: ENCODE [22, 32], Najafabadi et al. [33], Schmitges et al. [20], and Imbeault et al. [30]. ENCODE data were downloaded from ENCODE project website (https://www.encodeproject.org/experiments/), while the other data were downloaded from GEO (accession numbers GSE58341, GSE76494, and GSE78099). A total of 2209 ChIP-seq experiments were analyzed, covering 604 TFs and six cell lines (**Supplementary Table 1**).

Raw reads were aligned to the human reference genome (GRCh38) with *bowtie2* (version 2.3.4.1) using the *“--very-sensitive-local”* mode. Mapped reads with mapping quality score smaller than 30 were removed using *samtools* (version 1.9)[34]. ChIP-seq peaks were called using *MACS* (version 1.4) [35, 36] with a permissive p-value threshold of 0.01. We used this permissive p-value to obtain a range of TF binding signals, which our method uses to quantitatively model TF binding strength. We also included negative peaks, i.e. peaks obtained by swapping the treatment with the control experiments, to enable proper modeling of the background signal. In the end, for each ChIP-seq experiment, this process resulted in a list of peaks covering a wide range of pulldown or control (background) signal strengths, along with their associated read counts. The complete set of uniformly processed peaks used in this study can be accessed via Zenodo (DOI: 10.5281/zenodo.5573261).

### WGBS data processing and DNase-seq data retrieval

Raw reads from Whole-Genome Bisulfite Sequencing (WGBS) of six cell lines were retrieved from ENCODE and GEO (see **Supplementary Table 2** for accession numbers). Raw reads were trimmed based on their quality (phred33 ≥ 20) with *TrimGalore* (version 0.6.4) [37]. Paired reads were aligned to the human reference genome hg38 [38] using *bismark (bowtie2* mode, version 0.22.2), allowing one mismatch during alignment. Reads were deduplicated by removing those that aligned to the same genomic position (*bismark:deduplicate_bismark*). Methylation calls were then extracted, ignoring the first 2 bps from the 5’ end of read 2 (*bismark:bismark_methylation_extractor*). A genome-wide coverage report with methylated and unmethylated read counts was then generated (*bismark:coverage2cytosine*). Finally, a bigwig file was generated for unmethylated and methylated counts (*bedGraphToBigWig*)[39].

For DNase-seq data, read depth-normalized bigwig files representing DNase-seq signal were retrieved from ENCODE (see **Supplementary Table 2** for accession numbers).

### Formatting and preprocessing of data for JAMS

To retrieve the sequence, DNA accessibility, and DNA methylation to train our model, we focused on the positive and negative ChIP-seq peak regions that did not fall within endogenous repeat elements, since the homology of repeat elements can confound the modeling of ChIP-seq data based on sequence [33]. This was done by removing peaks that overlapped any repeat regions, as defined by RepeatMasker [38, 40].

To model the effect of sequence and epigenetic factors on TF binding using our method, it is necessary to align the peaks based on the position of the most likely TF binding site. To do so, we used the known motif of each TF, in the form of position frequency matrices (PFMs), to search for the most likely TFBS within the 100 bp range of the peak summit. PFMs were obtained from CIS-BP [25], and were augmented by *de novo* motifs identified by RCADE2 [41, 42] for the C2H2-ZF family of TFs as described in later sections. CISP-BP contains more than one PFMs per TF, as they are derived from different experimental techniques. We selected PFMs exclusively derived from *in vitro* experiments, in order to avoid the confounding effects present *in vivo*. We prioritized, in descending order, PFMs from SELEX, Selective microfluidics-based ligand enrichment followed by sequencing (SMiLE-seq), and Protein-Binding Microarrays (PBM). We used *AffiMx* [43] to identify the best motif match in each peak sequence. This process was uniformly applied to all peaks, including the negative ChIP-seq peak set.

Once the best motif hit in each peak was identified, we extracted the sequence and nucleotide-resolution methylation profile at the motif hit as well as the flanking regions (20 bp) around the motif hit. Sequences were retrieved from the reference genome hg38 using *bedtools:getfasta* [38, 44]. Methylated and unmethylated read counts at each position were retrieved from the WGBS bigwig files using *bwtool* [45].

Similarly, normalized DNA accessibility was extracted from the motif hit region and 500 bp upstream and downstream of the motif hit from the DNase-seq bigwig files. ChIP-seq read counts were extracted from the control and pull-down experiments for the +/− 400bp region surrounding the motif match using *bedtools:multicov* (MAPQ score > 30). (**Fig. 4C, bottom**) [44].

We emphasize that while a known motif of each TF was used to identify an offset for each peak and align the peak regions, this process is not expected to confound the sequence features learned by JAMS, since it is uniformly applied to all peaks regardless of the signal strength. The TF motifs themselves were also not used by JAMS, and the sequence features that are predictive of ChIP-seq signal were learned *de novo* from the aligned peaks.

### Implementation of JAMS

Our method creates a joint accessibility-methylation-sequence model (JAMS model) for each ChIP-seq experiment, in which the ChIP-seq signal of each peak is explained as a function of accessibility, methylation, and sequence at that peak. Consider the *k×m* matrix ***X***, which represents the value of *m* predictive features at *k* genomic positions (i.e. peaks). These *m* features include those related to accessibility (A), methylation (M), and sequence (S):

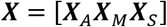

JAMS models the logarithm of TF binding strength at each of the *k* peaks as a linear function of the matrix ***X***:

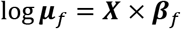

Here, ***μ***_*f*_ is the vector of the binding strength for transcription factor *f* across *k* peaks, ***X*** is the *k×m* feature matrix described above, and ***β***_*f*_ is the vector of *m* coefficients that describe the effect of each of the *m* features on the TF binding strength (matrices are denoted with bold capital letters, and vectors with bold lower-case letters).

Similarly, the background ChIP-seq signal across the peaks is also modeled as a function of ***X***:

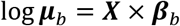

Here, ***μ***_*b*_ represents the background signal strength across *k* peaks, and ***β***_*b*_ is the vector of *m* coefficients that describe the effect of each of the *m* features on the background signal.

In a ChIP-seq experiment, the expected control (background) read counts at each peak is a function of the background signal multiplied by the library size. Therefore, the logarithm of control reads can be modeled as:

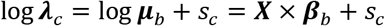

Here, ***λ***_*c*_ is the vector of expected (average) control read counts across the *k* peaks, and *s_c_* is an experiment-specific size factor that can be interpreted as the logarithm of sequencing depth for the control library.

The expected pull-down read counts in a ChIP-seq experiment, however, are a function of both the background signal and the TF binding strength, multiplied by the library size. Therefore:

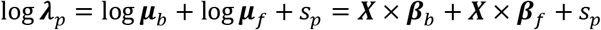

Here, ***λ***_*p*_ is the vector of expected pulldown read counts across the *k* peaks, and *s_p_* can be interpreted as the logarithm of sequencing depth for the pulldown library.

While these equations describe the expected control and pulldown read counts, the actual observed read counts are probabilistic observations that may deviate from these expected values. Here, we model the read counts as observations from negative binomial distributions [46] whose mean is given by the equations above, with a shared dispersion parameter across the peaks:

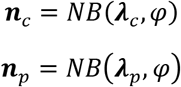

Here, ***n***_*c*_ and ***n***_*p*_ are the vectors of observed control and pulldown read counts across the *k* peaks, respectively, and *φ* is the dispersion parameter. The equations above allow us to jointly model the control and pulldown experiments as a function of ***X***. We use the glm.nb function in R for this purpose and fit a model of the form *n~XX+t+XX:t*, where *n* is an R vector that concatenates the observed control and pulldown read counts (with length 2*k*), *XX* is the result of duplicating matrix ***X***, i.e. *XX=rbind(X,X)*, and *t* is a binary vector of length 2*k* indicating whether the observed read count comes from the control experiment (0) or from the pulldown experiment (1). The coefficients returned by the glm.nb function for *XX* correspond to ***β***_*b*_ in the equations above, and the coefficients for *XX:t* correspond to ***β**_f_*. The glm.nb also returns the standard error of mean and a p-value for each of these coefficients, which we use to determine the statistical significance.

Constructing the matrix ***X***: Sequence, DNA methylation and DNA accessibility are used as the predictor variables, which are included in the matrix ***X***. We used one-hot encoding for the sequence over the TFBS. Methylated and unmethylated read counts over the motif were used to calculate the methylation percentage at each position. If the average coverage of methylation and unmethylated reads over the motif is less than 10 counts, the peak is removed. Average DNA accessibility was calculated for bins of 100 bp (10 bins) plus one bin for the TFBS region itself, and then logarithm of DNA accessibility was calculated; a pseudocount equivalent of 1% of the smallest value was used to allow for log transformation of the data. Average methylation percentage and sequence composition of the flanking regions were also used as predictors.

The source code for JAMS, as well as the complete set of JAMS models generated in this study are available at https://github.com/csglab/JAMS. Additional data, including the JAMS motif logos and the data used to train the JAMS models are deposited to Zenodo (DOI: 10.5281/zenodo.5573261).

### Differential binding analysis

To calculate differential TF binding between cell lines, we first identified CTCF ChIP-seq experiments from ENCODE that had at least two biological replicates per cell line (**Supplementary Table 5**), and retrieved the pull-down and control experiment data. After aligning and peak calling, we defined a unified list of peaks that were present in at least one sample. Peaks that were present in more than one sample and had summits within 100 bp of each other were merged, as they likely represent the same CTCF binding site. Then, the best motif match within 100 bp of each summit was identified [43]. We extracted ChIP-seq read counts present within a 400bp range from the motif hit in the pull-down and control experiments and created a count matrix.

We used DESeq2 [47] to compare the pulldown-to-control ratio between pairs of cell lines, limiting to comparisons that included only data from the same lab. The DESeqDataSetFromMatrix function from DESeq2 was used to create a DESeqDataSet object, followed by fitting a model of the form *~s*+*c: t*, where *s* is a categorical variable representing the sample/replicate (shared between pairs of control and pulldown experiments), *c* is a binary variable representing the two different cell lines, and *t* is a binary variable denoting whether the read count corresponds to the control experiment (0) or the pulldown experiment (1). After fitting the DESeq2 model, the coefficient for *c*:*t* corresponds to the log2 fold changes. Significant differentially bound peaks (FDR < 0.1) were identified for every pair of cell lines, excluding cell line pairs whose ChIP-seq experiments were done in different laboratories. The pair of cell lines (GM12878 and HeLa-S3) with the highest number of significantly bound peaks were selected for further analysis.

### Inference of PFMs for C2H2-ZF proteins using RCADE2

We inferred position frequency matrices (PFMs) for canonical C2H2 zinc finger proteins using RCADE2 [41, 42]. RCADE2 uses the protein sequence, the DNA sequence of the ChIP-seq peaks, and a previously computed machine learning-based recognition code to predict the DNA-binding preferences of C2H2-ZFPs. The protein sequences for these TFs were retrieved from UniProt [48]. We focused on the top 500 ChIP-seq peaks (sorted by p-value) that did not fall within endogenous repeat elements (EREs) [38, 40]. The DNA sequence of the +/− 250 region around the peak summits for the top 500 non-ERE peaks along with the protein sequence was provided as input to RCADE2, and the optimized motif was used to augment the CIS-BP motifs.

## ADDITIONAL FILES

**Additional file 1:** PDF file (.pdf) including Supplementary Figures.

**Additional file 2:** Excel file (.xlsx) including Supplementary Tables.

## DECLARATIONS

### Ethics approval and consent to participate

Not applicable.

### Consent for publication

Not applicable.

### Availability of data and materials

The source code for the method presented in this study, JAMS, is available via GitHub at https://github.com/csglab/JAMS. This GitHub repository also contains the models generated by JAMS as part of this study. Additional data, including the uniformly processed peaks and their tag counts, which were used to train the JAMS models, along with additional JAMS output files are available via Zenodo (DOI: 10.5281/zenodo.5573261).

### Competing interests

The authors declare that they have no competing interests.

### Funding

This work was supported by funds from the Natural Sciences and Engineering Research Council of Canada (RGPIN-2018-05962) and resource allocations from Compute Canada to HSN. AHC was partially supported by the Globalink Graduate Fellowship from Mitacs. HSN holds a Canada Research Chair funded by the Canadian Institutes of Health Research.

### Authors’ contributions

AHC and HSN developed the computational methods. AHC analyzed the data. AHC and HSN prepared the manuscript. HSN directed the study.

## Acknowledgements

We thank Senthilkumar Kailasam for assisting with analysis of WGBS data.

## Notes

https://github.com/csglab/JAMS

http://zenodo.org/record/5573261

### Competing Interest Statement

The authors have declared no competing interest.

