## Additional file 1: Supplementary Figures for "A base-resolution panorama of the *in vivo* impact of cytosine methylation on transcription factor binding"

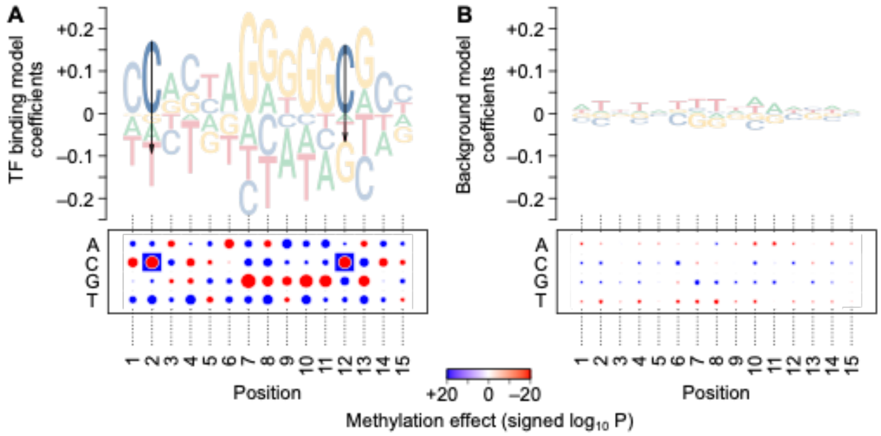


**Supplementary Figure 1. TF-specific and background coefficients for CTCF in HEK293 cells.** (**A**) Motif logo and dot plot representations of the sequence/methylation preference of the TF-specific signal. The logo (top) shows methylation coefficients as arrows, with the arrow length proportional to the mean estimate of methylation effect. The heatmap (bottom) shows the magnitude of the preference for each nucleotide at each position using the size of the dots, with red and blue representing positive and negative coefficients, respectively. The signed logarithm of P-value of the methylation coefficient is shown using the color of the squares around the dots, with red and blue corresponding to increased or decreased binding to methylated CpG, respectively (only significant methylation coefficients at FDR<1×10–5 are shown). Note that while the methylation coefficient corresponds to the entire CpG dinucleotide, only the C in the CpG dinucleotide is marked with the colored squares. (**B**) Motif logo and dot plot representations for the background signal.

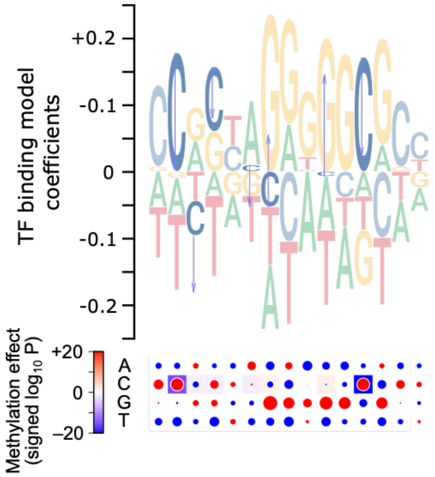


**Supplementary Figure 2. Likelihood ratio test per position to identify CTCF binding site positions with significant methylation effects.** For each position of the binding site, a reduced model was trained, each exclusing methylation of that position from the predictive variables. Then, each of these reduced models were compared to the whole CTCF JAMS model using a likelihood ratio test (LRT). The p-values obtained from the LRT are shown as the color of the squares. The effect sizes for the bases and methylation are obtained from the full CTCF JAMS model. Significant LRT p-values indicate that removing the methylation of the corresponding position from the model reduces the goodness of fit. The motif logo and dot plot representations follow the same notations as **Supplementary Fig. 1**.

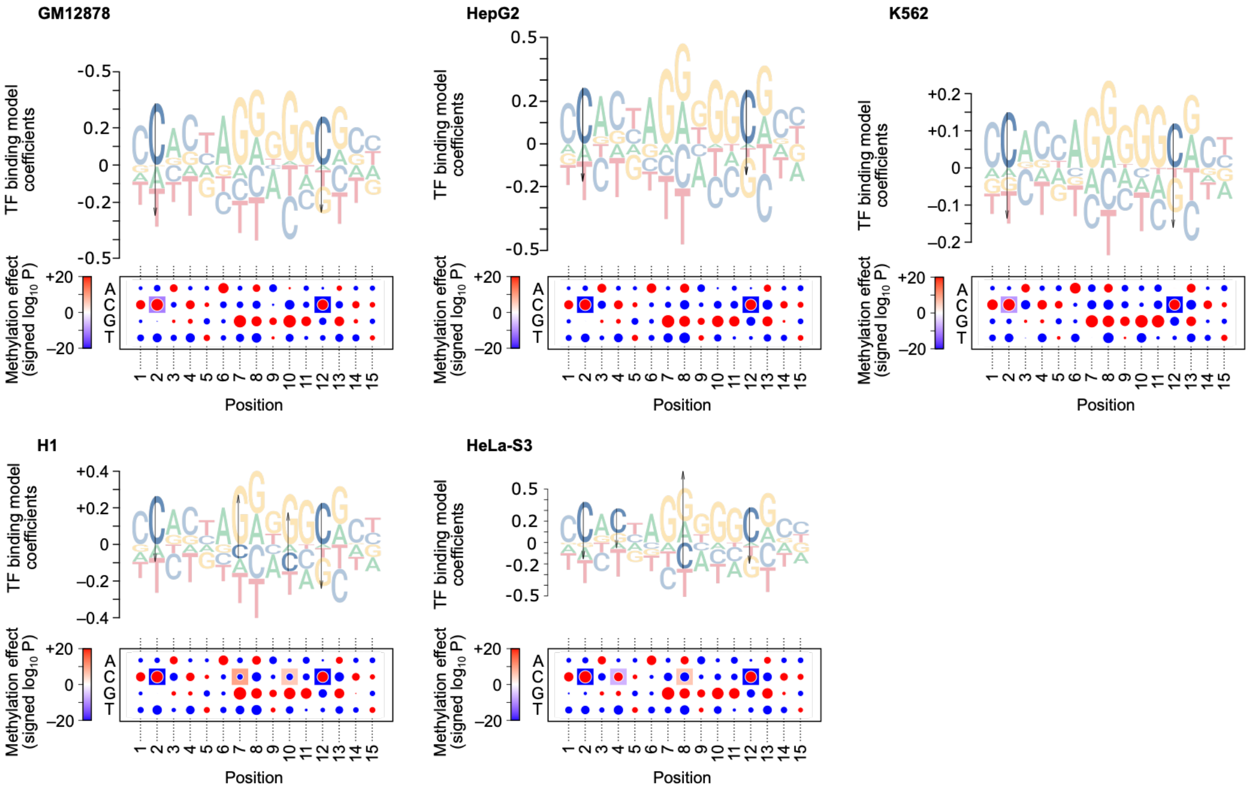


**Supplementary Figure 3. JAMS coefficients for CTCF across different cell lines.** Motif logs and dot plot representations follow the same format as described in **Supplementary Fig. 1**. TF-specific and background model coefficients are shown side-by-side for each of the six cell lines (GM12878, HepG2, H1, HeLa-S3, and K562).

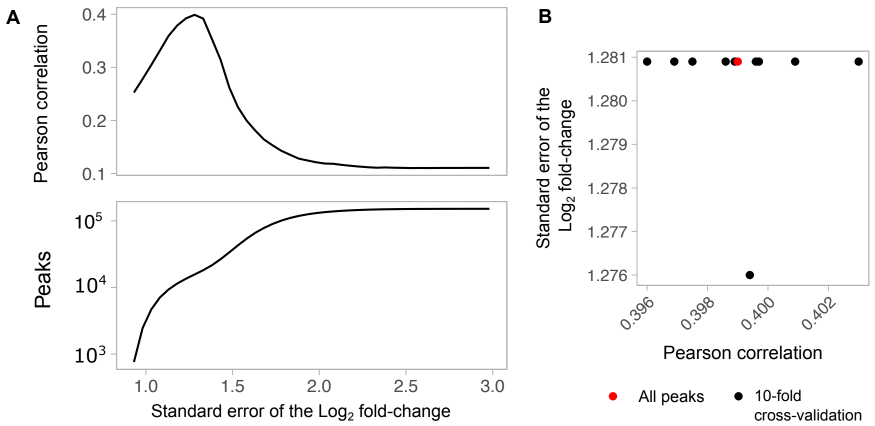


**Supplementary Figure 4. Calculating logFC S.E.M. threshold.** (**A**) Pearson correlation between the predicted and observed change in CTCF binding, after filtering the CTCF peaks based on different cutoffs for standard error of mean (S.E.M.) of the LFC of pull-down/control ratio. An optimal threshold is observed at logFC S.E.M. = 1.28. (**B**) To discard the possibility of overfitting of the threshold, different optimal thresholds were calculated using a 10-fold cross-validation approach. Specifically, each time 90% of the peaks were used to identify the optimal logFC S.E.M threshold, and the Pearson correlation between the predicted and observed peaks that passed that threshold was calculated on the remaining 10% of peaks The logFC S.E.M. threshold obtained by using all peaks (red dot) is similar to the thresholds obtained with cross-validation (black dots), and leads to a similar correlation between the predicted and observed change in CTCF binding for the held-out peak sets (ranging from 0.396 to 0.403).

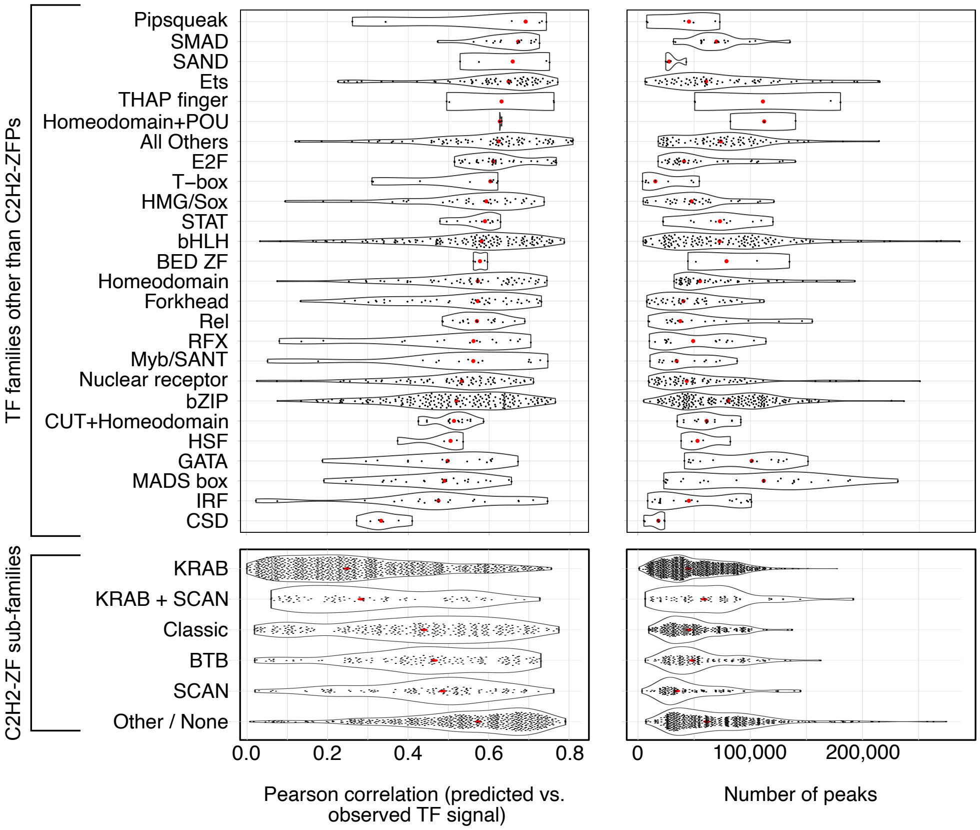


**Supplementary Figure 5. MethylChIP results by TF families.** Violin plots showing the Pearson correlation between observed and predicted pull-down tag density (left) and number of peaks used to train the GLM (right), shown separately for each TF family (top) and C2H2-ZF subfamilies (bottom).

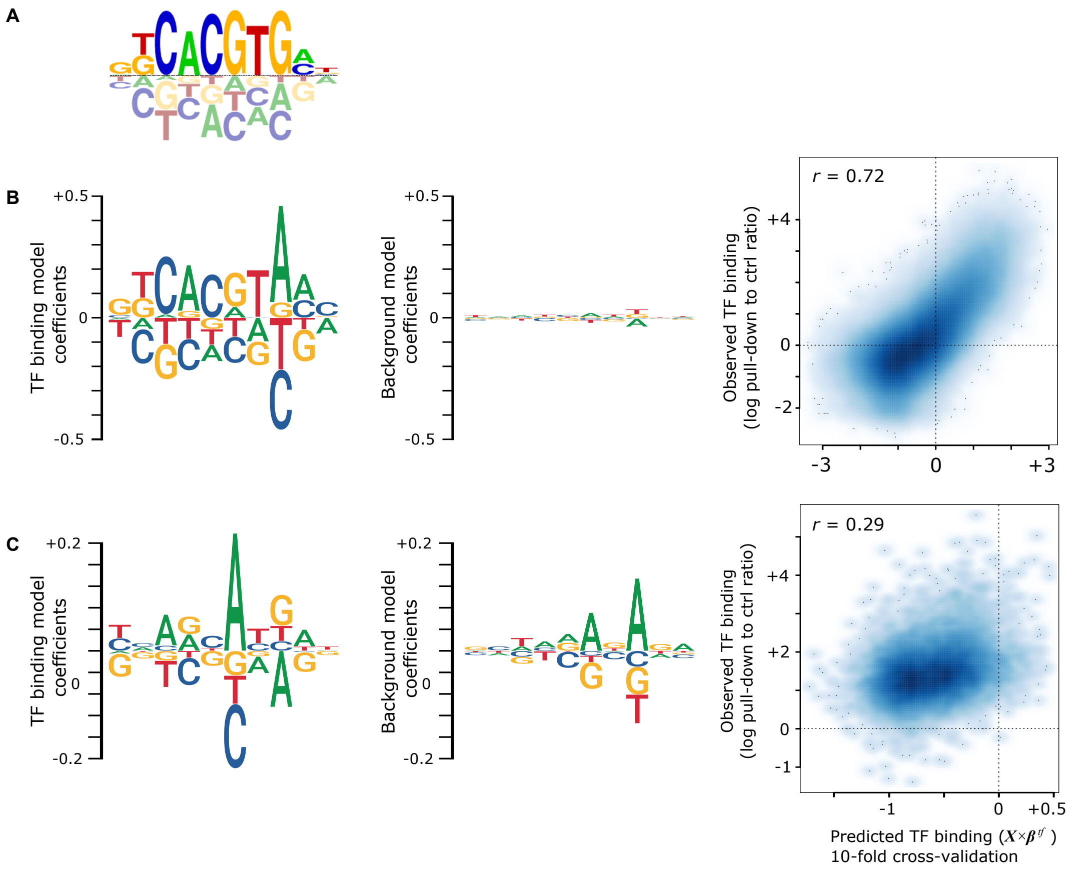


**Supplementary Figure 6. Example high-quality and low-quality JAMS models.** (**A**) The known BHLHE40 motif, obtained from the CIS-BP database, shown as an example (motif ID M02788_2.00). (**B-C**) Results from a high-quality (**B**) and a low-quality (**C**) JAMS model for BHLHE40. Inferred sequence coefficients for TF binding (left) and background (middle), as well as the predicted vs. observed TF binding signal (right) are shown.

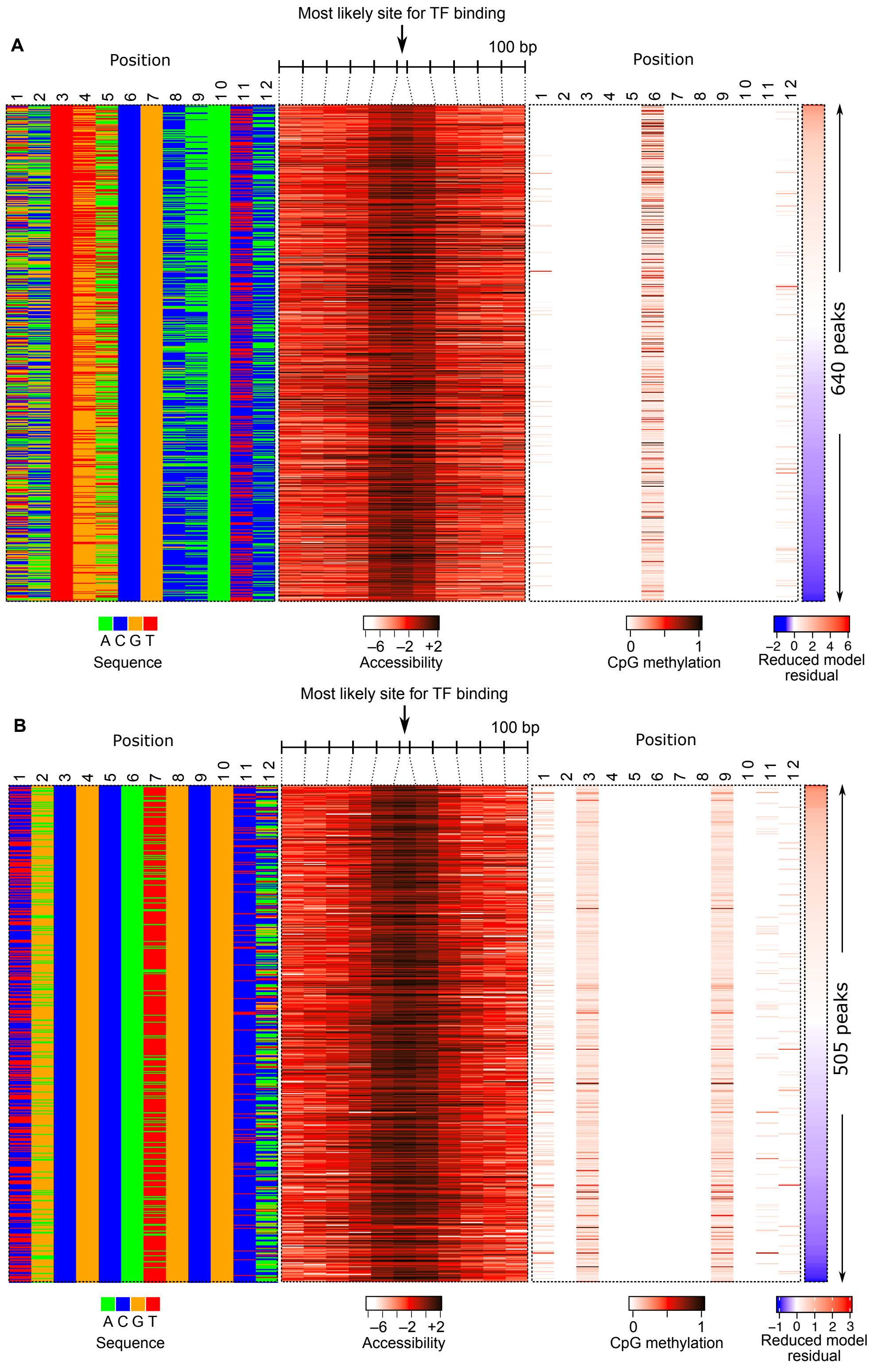


**Supplementary Figure 7. *In vivo* methylation binding preferences of CEBBP and NFR1.** (**A**) Heatmap representation of the sequence, accessibility, and CpG methylation, for a subset of CEBPB peaks that have high DNA accessibility, are similar to the CEBPB consensus binding sequence, and have a CpG in position 6/7. Peaks are sorted by the residual of a reduced JAMS model that does not use the methylation level for predicting the TF binding signal. (**B**) Same as panel A, but for NFR1 (with the requirement to have CpG dinucleotides at positions 3/4 and 9/10).

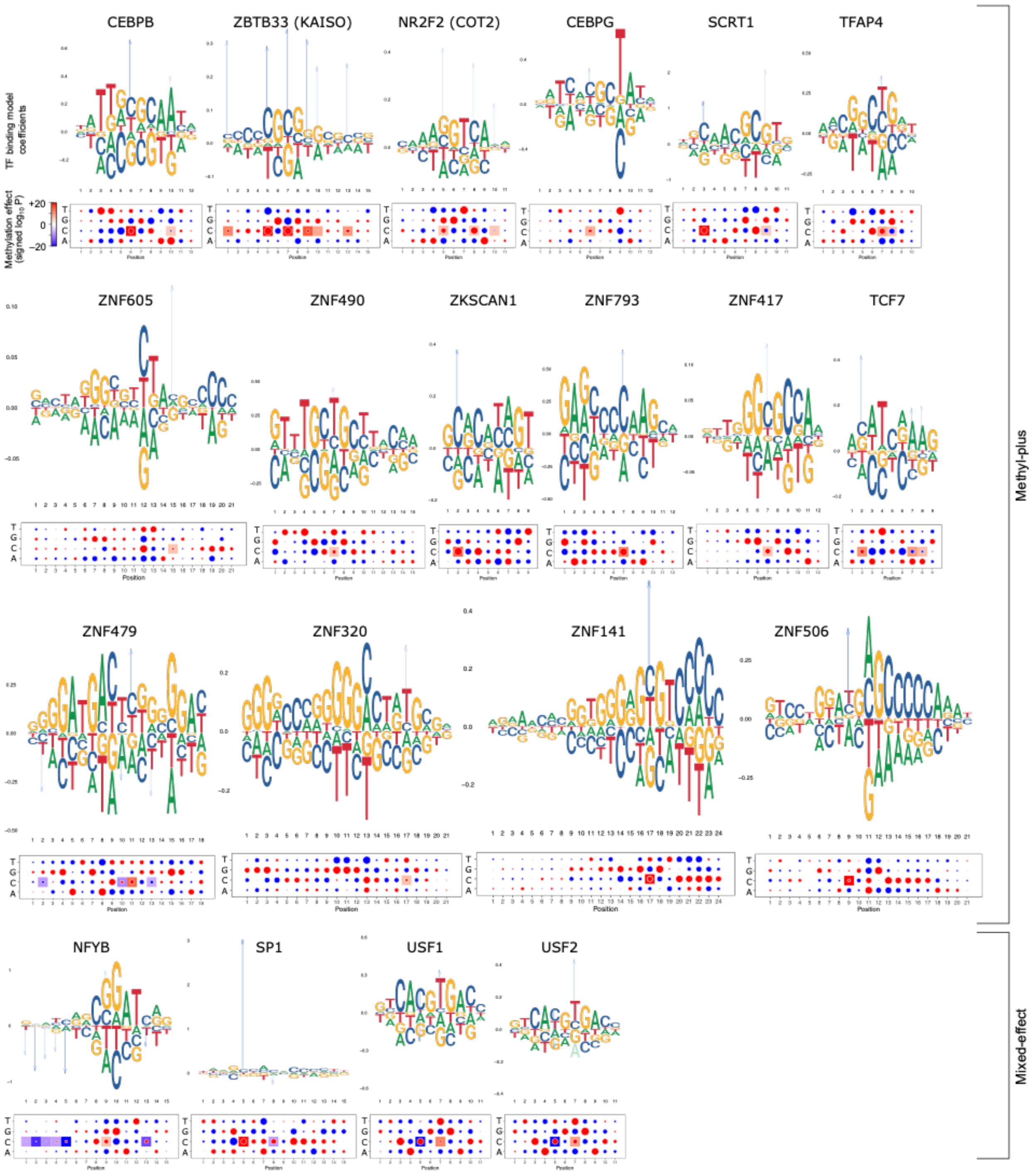


**Supplementary Figure 8. Methyl-plus and mixed-effect TFs identified by JAMS.** Motif logo and dot plot representations follow the same formatting as **Supplementary Fig. 1**.
